# Spatial cholesterol homeostasis gatekeeps T-cell development and activation by orchestrating signaling and fitness

**DOI:** 10.64898/2026.08.16.745088

**Authors:** Yanchi Li, Xing He, Chenxi Li, Zhengxu Ren, Jing Huang, Qian Yang, Mingming Gao, Yingjie Wu, Xiwei Liu, Chenqi Xu

**Author notes:** These authors contributed equally (Yanchi Li, Xing He, Chenxi Li). These authors jointly supervised this work (Xiwei Liu, Chenqi Xu).

## Abstract

Cholesterol is essential for T-cell immunity, and its spatial distribution is tightly regulated. Although cholesterol is synthesized in the endoplasmic reticulum (ER), it is predominantly transported to the plasma membrane (PM); however, the machinery mediating this anterograde transport in T cells remains unknown. Here, through a functional genetic screen, we identify oxysterol-binding protein (OSBP) as the principal mediator of ER-to-PM cholesterol transport in T cells. OSBP deficiency depletes accessible PM cholesterol while causing cholesterol accumulation in the ER, resulting in impaired T-cell receptor (TCR) signaling and disruption of ER homeostasis. Using stage-specific conditional knockout mice, we demonstrate that OSBP is required at multiple developmental checkpoints in the thymus, including β-selection, positive selection, and post-selection maturation. Loss of OSBP during early thymocyte development causes a near-complete block in T-cell development, resulting in a profound absence of mature peripheral T cells. In mature T cells, activation markedly increases dependence on OSBP-mediated cholesterol transport, with its inhibition causing ER perturbation and extensive cell death. Finally, we show that disease-associated oxysterols disrupt OSBP-mediated cholesterol transport, leading to T-cell dysfunction and providing a mechanistic explanation for impaired T-cell immunity in pathological settings. Together, our findings identify OSBP as a central regulator of intracellular cholesterol transport that couples membrane cholesterol homeostasis to TCR signaling and ER integrity. These results establish the spatial distribution of cholesterol, rather than its abundance alone, as a fundamental metabolic determinant of thymocyte development and peripheral T-cell function.

## INTRODUCTION

Cholesterol is an essential structural component of cellular membranes and a critical regulator of membrane organization and signal transduction^1^. To meet the increased demand for membrane biogenesis during proliferation, activated T cells upregulate cholesterol biosynthesis and uptake^2^. Cholesterol homeostasis also critically shapes the differentiation and function of diverse T-cell subsets, including cytotoxic T cells, Th1, Th2, Th17, Tfh, and Treg cells^3–11^. Dysregulation of this balance can consequently compromise T-cell function across different pathological settings. Within the tumor microenvironment, tumor-derived oxysterols can induce cholesterol deficiency in T cells and thereby dampen antitumor immunity^3^. Conversely, increasing T-cell cholesterol levels by inhibiting the cholesterol esterification enzyme ACAT1 (SOAT1 in humans) improves the efficacy of immune checkpoint blockade, chemotherapy, cancer vaccination, and adoptive T-cell therapy^12–15^. Serum cholesterol levels, particularly low-density lipoprotein (LDL) cholesterol levels, are positively correlated with CAR-T cell expansion in patients^16^. In autoimmune settings, enhanced cholesterol biosynthesis in CD4⁺ T cells drives pathogenic Th1 differentiation in inflammatory bowel disease (IBD)^4^, whereas statin use has been associated with a reduced risk of IBD^17^. Collectively, these studies establish cholesterol homeostasis as a fundamental determinant of T-cell immunity.

Cholesterol is synthesized primarily in the endoplasmic reticulum (ER) and subsequently redistributed across cellular membranes, with the highest abundance at the plasma membrane (PM)^18^. PM cholesterol is partitioned into three functionally distinct pools: an accessible (active) pool, a sphingomyelin-sequestered pool, and an essential (inaccessible) pool^19^. Among these, accessible cholesterol, which can be detected using the ALOD4 probe, is particularly dynamic and responsive to changes in cellular state^3,20^. Upon T-cell activation, the accessible PM cholesterol pool increases markedly^21^. Cholesterol directly interacts with the transmembrane region of the TCR-CD3 complex, promoting TCR clustering while contributing to the maintenance of the resting TCR in an inactive conformation^12,22–24^. Cholesterol also increases membrane lipid packing, thereby weakening electrostatic interactions between the basic residue-rich sequences (BRSs) of CD3 chains and acidic phospholipids^25–27^. This facilitates the release of the BRSs and adjacent immunoreceptor tyrosine-based activation motifs (ITAMs) from the membrane, enabling their phosphorylation and downstream signaling. Thus, PM cholesterol organization is intimately coupled to TCR signaling. Beyond its direct effects on TCR signaling, PM cholesterol supports the organization of specialized membrane domains, including the immunological synapse. Signaling molecules are spatially organized into distinct domains to coordinate stimulatory and inhibitory signals and sustain T-cell responses^28^.

PM cholesterol homeostasis is tightly regulated by anterograde and retrograde transport. Non-vesicular lipid transfer proteins enable rapid lipid exchange between cellular membranes and are critical for establishing lipid gradients and maintaining membrane lipid homeostasis^29–31^. Aster-A has been identified as a key protein transporting cholesterol from the PM to the ER in T cells^21^. However, it is still unclear how cholesterol is transported from ER to PM in T cells. Oxysterol-binding protein (OSBP) and OSBP-related proteins (ORPs) constitute a highly conserved family of lipid transfer proteins involved in intracellular sterol transport and membrane lipid organization^32–34^. The mammalian ORP family comprises 12 members with distinct roles. However, the functions of the ORP family in T cells remain largely unexplored.

Here, we first performed a functional screen of the ORP family and identified OSBP as a key transporter that maintains PM cholesterol in T cells. We next examined the impact of OSBP perturbation on cholesterol distribution, TCR signaling, and ER homeostasis. Using stage-specific *Osbp* conditional knockout mice and *ex vivo* perturbation, we examined the requirement for this pathway across T-cell developmental stages and activation states. We further extended these analyses to human T cells and assessed how disease-associated oxysterols influence OSBP-regulated cholesterol distribution and T-cell function.

## RESULTS

### OSBP mediates intracellular cholesterol transport toward the plasma membrane in T cells

Given the crucial roles of cholesterol in T cells, we first characterized cholesterol levels across distinct T-cell developmental stages. Accessible PM cholesterol was quantified using the cholesterol-binding probe ALOD4, whereas total cellular cholesterol was measured by Filipin III staining. Among thymocyte subsets, double-negative (DN) thymocytes exhibited substantially higher levels of accessible PM cholesterol compared with double-positive (DP) and single-positive (SP) thymocytes (Fig. 1A). Consistently, total cellular cholesterol followed a similar pattern across thymocyte subsets, with DN thymocytes showing the highest levels (Fig. 1B). Peripheral mature CD4⁺ and CD8⁺ T cells generally displayed lower levels of both accessible PM cholesterol and total cellular cholesterol than thymocytes, whereas accessible PM cholesterol levels were comparable to those observed in DP thymocytes (Figs. 1A, B). These findings indicate that PM cholesterol levels are dynamically regulated during T-cell development, highlighting the need for a better understanding of intracellular cholesterol transport.

**Figure 1.**
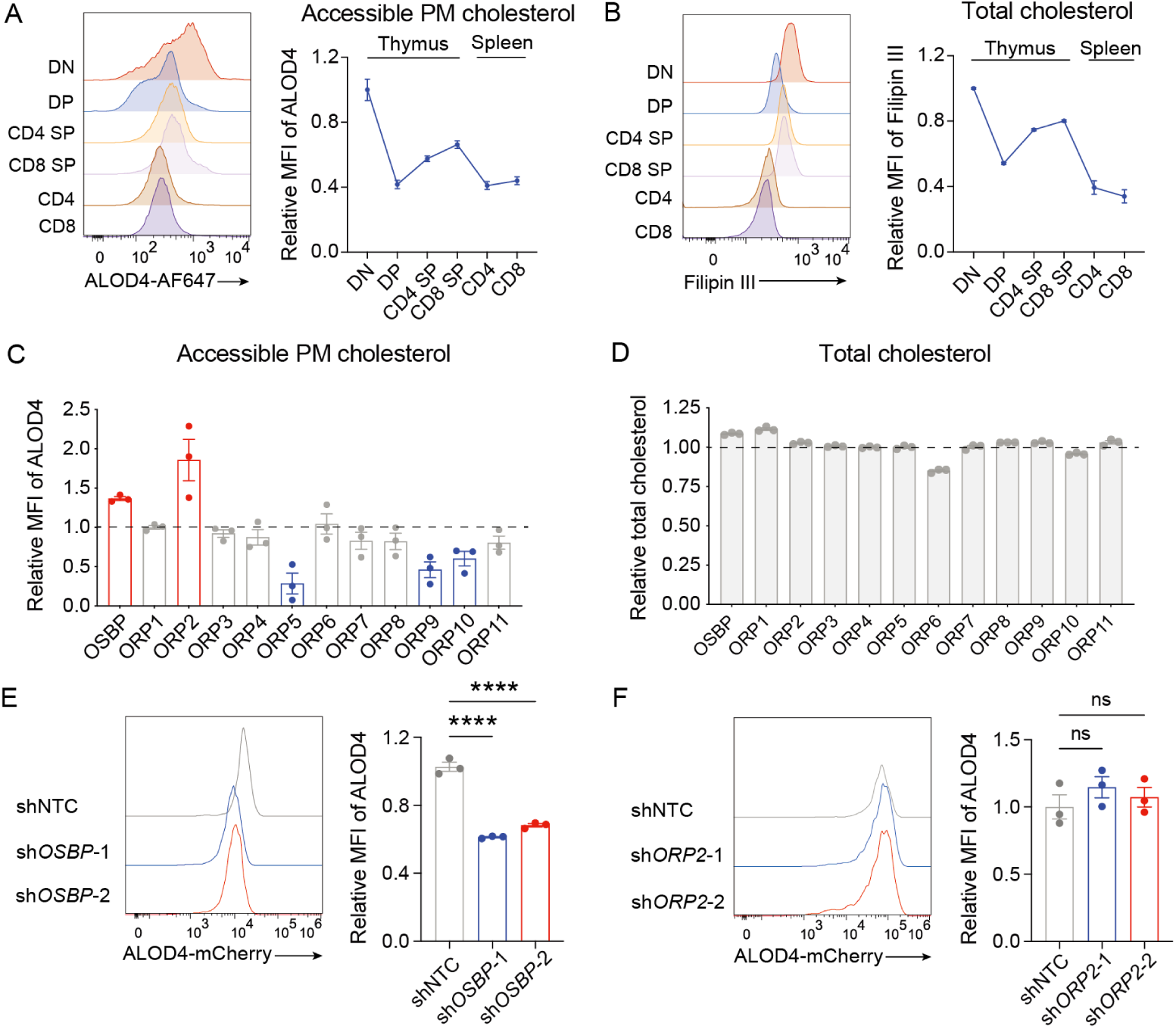
Identification of OSBP as a key transporter maintaining plasma membrane cholesterol availability in T cells. (A-B) Accessible plasma membrane (PM) cholesterol levels (A) and total cellular cholesterol levels (B) in double-negative (DN), double-positive (DP), CD4 single-positive (CD4 SP), and CD8 single-positive (CD8 SP) thymocytes, as well as peripheral CD4⁺ and CD8⁺ T cells, measured using ALOD4 and Filipin III staining, respectively. Representative flow cytometry profiles and quantification of median fluorescence intensity (MFI) normalized to the corresponding values in DN thymocytes are shown (n = 4 mice). (C-D) Functional screen of ORP family members in Jurkat T cells. Accessible PM cholesterol levels (C) and total cellular cholesterol levels (D) in cells ectopically expressing individual ORP family members were measured using ALOD4 staining and an Amplex Red cholesterol assay, respectively. Values were normalized to the corresponding batch-matched vector controls (n = 3). (E-F) Effects of *OSBP* or *ORP2* knockdown on accessible PM cholesterol levels in Jurkat T cells. Cells transduced with independent shRNAs targeting *OSBP* (E) or *ORP2* (F), together with non-targeting shRNA controls, were analyzed by ALOD4 staining. Representative flow cytometry profiles and quantification of ALOD4 MFI, normalized to non-targeting shRNA controls, are shown (n = 3). Data are presented as mean ± s.e.m. Statistical significance was assessed by one-way ANOVA (E, F). ns, not significant; ****p < 0.0001.

To identify candidate transporters responsible for anterograde cholesterol transport in T cells, we performed a functional screen of all 12 mammalian ORPs. Each ORP family member was individually overexpressed in Jurkat T cells, followed by assessment of accessible PM cholesterol levels using ALOD4 staining. Only OSBP and ORP2 markedly increased PM cholesterol levels, whereas the remaining ORPs exhibited minimal effects or reduced PM cholesterol levels (Fig. 1C). Notably, ORP overexpression caused only modest changes in total cellular cholesterol abundance (Fig. 1D), indicating that these proteins primarily regulate intracellular cholesterol distribution rather than overall cholesterol levels.

To further determine which candidate was required for maintaining PM cholesterol abundance, we performed loss-of-function analyses. Knockdown of *OSBP*, but not *ORP2*, significantly reduced accessible PM cholesterol levels (Figs. 1E, F; Fig. S1).

Together, these results identify OSBP and ORP2 as transporters mediating anterograde cholesterol transport in T cells, while only OSBP is indispensable.

### OSBP orchestrates T cell signaling and ER fitness

To investigate the functional roles of OSBP in T cells, we pharmacologically inhibited OSBP with OSW-1, a selective and potent inhibitor^35^. OSBP inhibition markedly reduced accessible PM cholesterol levels (Fig. 2A) and impaired TCR signaling, as demonstrated by decreased phosphorylation of key signaling molecules, including CD3ζ, ZAP70, PLCγ1, and ERK following TCR stimulation (Fig. 2B). OSBP inhibition also resulted in reduced cell numbers and increased phosphatidylserine exposure, as assessed by Annexin V staining (Figs. 2C, D). Importantly, ectopic expression of OSBP substantially alleviated the OSW-1-induced decrease in cell numbers (Fig. 2E), supporting the specificity of OSW-1-mediated effects. Although supplementation with exogenous cholesterol effectively restored accessible PM cholesterol levels in T cells treated with OSW-1, it failed to rescue the reduction in cell number, suggesting an alternative causal mechanism (Figs. 2A, C).

**Figure 2.**
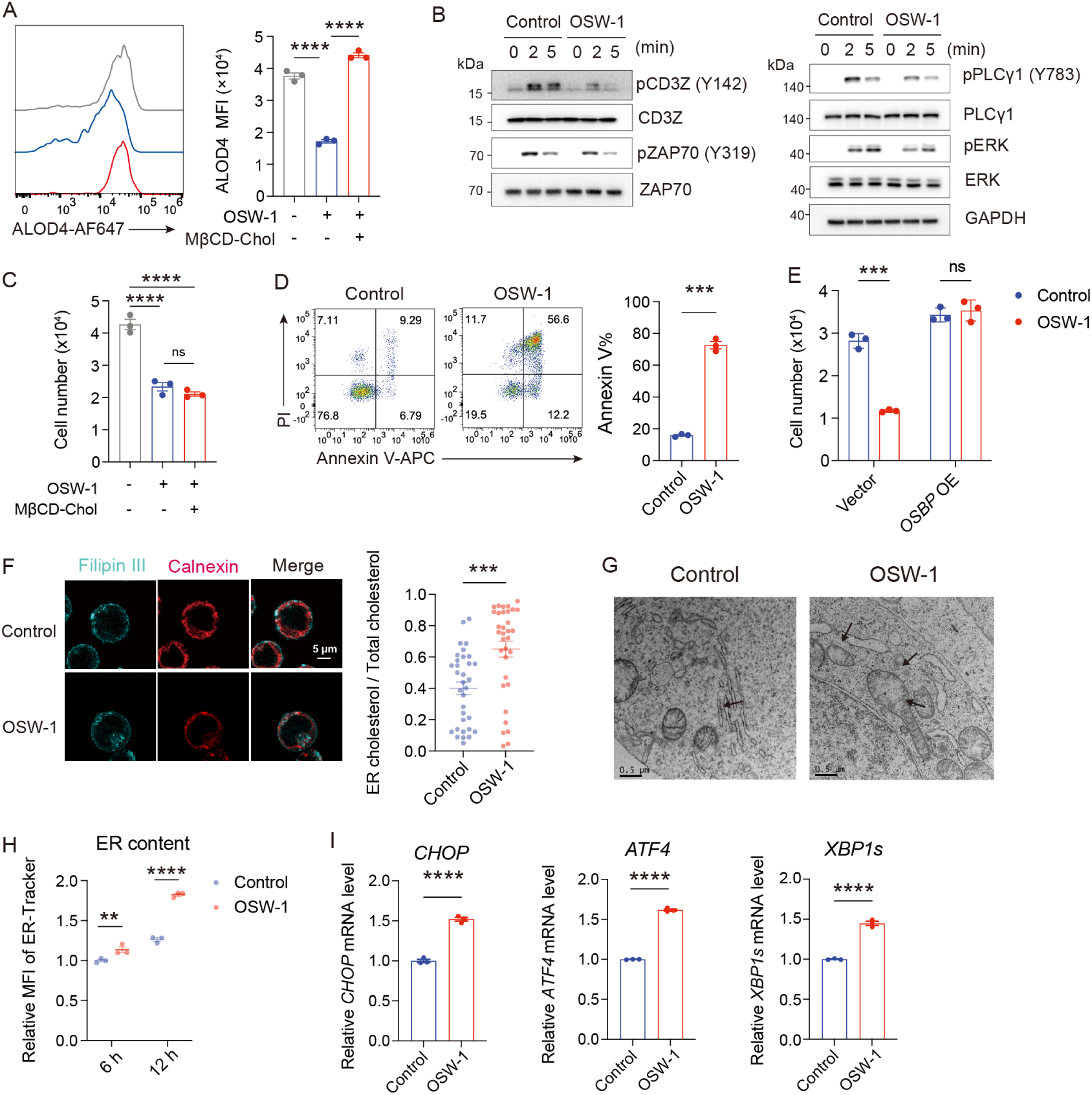
OSBP regulates cholesterol partitioning to maintain T-cell signaling and fitness. (A) Accessible PM cholesterol levels in Jurkat T cells following OSBP inhibition, measured by ALOD4 staining. Cells were treated with OSW-1 (5 nM, unless otherwise described) or DMSO control for 4 h. Where indicated, cells were co-treated with methyl-β-cyclodextrin (MβCD)-cholesterol (5 μg/mL) for 4 h (n = 3). (B) TCR signaling in Jurkat T cells after OSBP inhibition. Cells were pretreated with OSW-1 (50 nM) or DMSO control for 4 h and stimulated with anti-CD3 (10 μg/mL) for the indicated times. Phosphorylated and total CD3ζ, ZAP70, PLCγ1 and ERK were analyzed by immunoblotting. (C-D) Effects of OSBP inhibition on T-cell fitness. (C) Cell numbers were quantified after 20 h of OSW-1 treatment, with MβCD-cholesterol supplementation as indicated (n = 3). (D) Annexin V/PI staining profiles and quantification of Annexin V^+^ cells after OSW-1 treatment (n = 3). (E) Effects of *OSBP* overexpression on Jurkat T cell numbers following OSW-1 treatment. Cells were transduced with *OSBP* or control vectors and treated with OSW-1 or DMSO control. Cell numbers were quantified after 20 h (n = 3). (F) Subcellular cholesterol distribution in Jurkat T cells following OSBP inhibition. Cells were treated with OSW-1 or DMSO for 4 h, and cholesterol localization was visualized by Filipin III staining together with the ER marker calnexin. Representative confocal images and quantification of the fraction of Filipin III signal colocalized with calnexin, as determined by the thresholded Manders’ coefficient (tM1), are shown (n = 33 cells). Scale bar, 5 μm. (G) Representative transmission electron microscopy images of ER morphology in Jurkat T cells treated with OSW-1 or DMSO control. ER dilation and vacuolar structures are indicated by arrows. Scale bars, 500 nm. (H) ER content in Jurkat T cells following OSBP inhibition. Cells were treated with OSW-1 or DMSO control for the indicated times and analyzed by ER-Tracker staining. MFI values were normalized to the 6 h control group (n = 3). (I) Relative mRNA levels of ER stress-associated genes (*CHOP, ATF4* and *XBP1s*) in Jurkat T cells, assessed by quantitative PCR. Cells were treated with OSW-1 or DMSO control for 6 h, and expression levels were normalized to DMSO-treated controls (n = 3). Data are presented as mean ± s.e.m. Statistical significance was assessed by one-way ANOVA (A, C), unpaired two-tailed Student’s t test (D, F, I), multiple t tests (E), or two-way ANOVA (H). ns, not significant; **p < 0.01, ***p < 0.001, ****p < 0.0001.

Confocal microscopy revealed that OSBP inhibition caused a pronounced accumulation of cholesterol within the ER (Fig. 2F). Given that excessive ER cholesterol accumulation can perturb ER homeostasis^36^, we next examined ER morphology and stress responses. Transmission electron microscopy revealed extensive ER dilation and vacuolar structures in OSBP-inhibited cells (Fig. 2G). ER-Tracker staining showed that ER content was increased (Fig. 2H). Consistently, OSBP inhibition activated unfolded protein response (UPR) pathways, as evidenced by increased expression of *CHOP*, *ATF4*, and *XBP1s* (Fig. 2I).

Collectively, these findings show that OSBP functions as a critical regulator of intracellular cholesterol partitioning. OSBP inhibition caused membrane cholesterol dysregulation at both the PM and ER, dampening TCR signaling and ER fitness.

### OSBP is essential for thymocyte development

The dynamic changes in PM cholesterol during thymocyte development suggested that OSBP might have stage-specific functions in thymopoiesis. To investigate this possibility, we generated *Osbp^flox/flox^*(*Osbp^f/f^)* mice and crossed them with *Cd2^Cre^* or *Cd4^Cre^* mice, which mediate gene deletion at early and late stages of thymocyte development, respectively^37,38^.

Crossing *Osbp^f/f^* mice with *Cd2^Cre^*mice led to OSBP depletion beginning at the DN stage^37^. OSBP deficiency caused a dramatic reduction in total thymic cellularity (Fig. 3A), with DP thymocytes and both CD4 SP and CD8 SP thymocyte populations being largely absent (Fig. 3B). We next examined DN subsets based on CD44 and CD25 expression, as productive TCRβ rearrangement and pre-TCR-mediated β-selection occur at the DN3 stage. While the frequencies of DN1 and DN2 thymocytes were relatively preserved, the DN3-to-DN4 transition was nearly completely blocked (Fig. 3C). These findings demonstrate that OSBP is indispensable for progression through the DN3-to-DN4 β-selection checkpoint.

**Figure 3.**
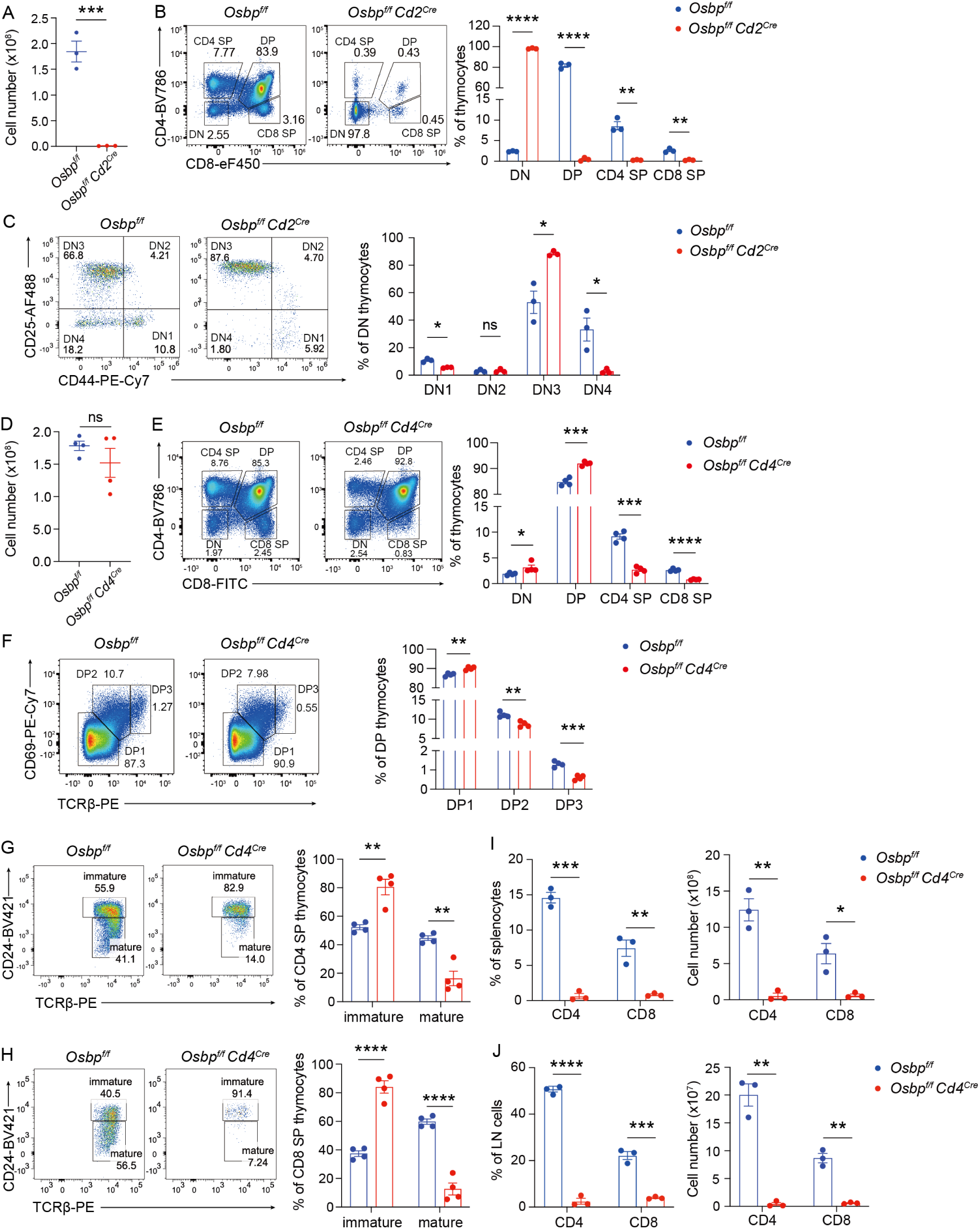
OSBP is required for thymocyte development at multiple developmental stages. (A-C) Thymocyte development in *Osbp^f/f^ Cd2^Cre^* mice and *Osbp^f/f^* control littermates (n = 3 mice). (A) Total thymic cell numbers. (B) Representative flow cytometry profiles and percentages of DN, DP, CD4 SP and CD8 SP cells among total thymocytes. (C) Characterization of DN thymocyte development. DN thymocytes were subdivided into DN1-DN4 subsets based on CD44 and CD25 expression, and subset frequencies within the DN population were quantified. (D-F) Thymocyte development in *Osbp^f/f^ Cd4^Cre^* mice and *Osbp^f/f^* control littermates (n = 4 mice). (D) Total thymic cell numbers. (E) Representative flow cytometry profiles and percentages of DN, DP, CD4 SP, and CD8 SP thymocytes among total thymocytes. (F) DP thymocyte progression through positive selection. DP thymocytes were subdivided into DP1-DP3 populations based on CD69 and TCRβ expression, and subset frequencies within the DP population were quantified. (G-H) SP thymocyte maturation in *Osbp^f/f^ Cd4^Cre^* mice and *Osbp^f/f^* control littermates (n = 4 mice). Frequencies of immature (TCRβ^+^CD24^+^) and mature (TCRβ^+^CD24^-^) cells within CD4 SP (G) and CD8 SP (H) thymocytes were quantified. (I-J) Peripheral T-cell populations in *Osbp^f/f^ Cd4^Cre^* mice and *Osbp^f/f^* control littermates (n = 3 mice). Frequencies and absolute numbers of CD4⁺ and CD8⁺ T cells in the spleen (I) and inguinal lymph nodes (J) were quantified. Data are presented as mean ± s.e.m. Statistical analysis was performed using unpaired two-tailed t test (A, D) or multiple unpaired two-tailed t tests (B, C, E-J). ns, not significant; *p < 0.05, **p < 0.01, ***p < 0.001, ****p < 0.0001.

To determine whether OSBP also regulates later stages of thymocyte development, we generated *Osbp^f/f^ Cd4^Cre^* mice, in which gene deletion begins at the DP stage. Although the reduction in total thymic cellularity was less severe than that observed in *Osbp^f/f^ Cd2^Cre^* mice (Fig. 3D), both CD4 SP and CD8 SP thymocyte populations were substantially reduced (Fig. 3E). We next subdivided DP thymocytes into DP1-DP3 subsets based on graded CD69 and TCRβ expression. TCRα rearrangement is initiated in DP1 thymocytes, whereas DP2 thymocytes represent the onset of positive selection following TCR recognition of self-peptide-MHC complexes^39,40^. OSBP deficiency resulted in a modest accumulation of DP1 cells accompanied by reductions in both DP2 and DP3 populations, with the latter being more profoundly affected (Fig. 3F), indicating impaired progression through positive selection. Furthermore, maturation of SP thymocytes was compromised in the absence of OSBP, with a more pronounced defect in CD8 SP than in CD4 SP cells (Figs. 3G, H). Consistent with these developmental defects, mature peripheral T cells were largely absent in the spleen and lymph nodes of OSBP-deficient mice (Figs. 3I, J).

To determine whether this function is shared by other members of the ORP family, we generated *Orp2^f/f^ Cd4^Cre^*mice. In contrast to OSBP deficiency, loss of ORP2 had no detectable effect on thymocyte development or peripheral T-cell homeostasis (Figs. S2A-C), further supporting OSBP as the principal ORP family member mediating anterograde cholesterol transport in T cells.

To further determine whether the developmental defects caused by OSBP deficiency depend on TCR repertoire diversity, we introduced the OT-I TCR transgene into *Osbp^f/f^ Cd4^Cre^* mice to generate a monoclonal TCR repertoire. Despite the restricted TCR specificity, OSBP-deficient OT-I mice exhibited reduced DP3 thymocytes and impaired generation of CD8 SP thymocytes (Figs. S3A, B). Moreover, the proportion of mature CD8 SP thymocytes was further decreased (Fig. S3C), indicating that OSBP is intrinsically required for thymocyte development independently of TCR repertoire diversity.

Together, these findings establish that OSBP is required at multiple developmental checkpoints during thymopoiesis, including β-selection, positive selection, and post-selection maturation.

### OSBP regulates distinct thymocyte developmental checkpoints through shared but stage-specific mechanisms

Our genetic models revealed that OSBP deficiency impaired multiple developmental checkpoints during thymocyte differentiation, including the DN3-to-DN4, DP2-to-DP3, and immature-to-mature SP transitions. Among these, the DN3-to-DN4 transition was most profoundly affected, resulting in an almost complete developmental blockade (Figs. 3B-C). Notably, these checkpoints depend on pre-TCR or TCR signaling. Unlike antigen-dependent TCR signaling, pre-TCR signaling occurs independently of antigen engagement and is primarily driven by spontaneous receptor clustering, a process that is thought to rely on plasma membrane cholesterol^41,42^.

Next, NUR77-GFP reporter mice were employed to assess pre-TCR/TCR signaling. After crossing NUR77-GFP mice with *Osbp^f/f^ Cd2^Cre^* mice, we analyzed NUR77 signal across DN1-DN4 subsets. NUR77-GFP expression was markedly reduced in DN4 thymocytes but remained largely unchanged in DN1-DN3 cells (Fig. 4A), indicating that OSBP is specifically required for pre-TCR signaling during β-selection. Furthermore, we crossed NUR77-GFP mice with *Osbp^f/f^ Cd4^Cre^* mice and analyzed DP and SP thymocytes. DP3 thymocytes exhibited substantially stronger NUR77-GFP signals than DP1 and DP2 cells, consistent with active positive selection, and OSBP deficiency significantly attenuated this signaling (Fig. 4B). In contrast, TCR signaling in immature CD4 SP and CD8 SP thymocytes was largely unaffected, whereas mature CD4 SP and CD8 SP thymocytes displayed moderately increased NUR77-GFP expression in the absence of OSBP (Figs. 4C, D). Notably, NUR77 signal progressively increased during thymocyte maturation, reaching its highest level in mature SP cells (Figs. 4A-D). The requirement for OSBP-mediated PM cholesterol transport was most pronounced at developmental stages characterized by relatively weak pre-TCR or TCR signaling, suggesting that cholesterol availability becomes a limiting factor when signaling strength is close to the threshold required for developmental progression.

**Figure 4.**
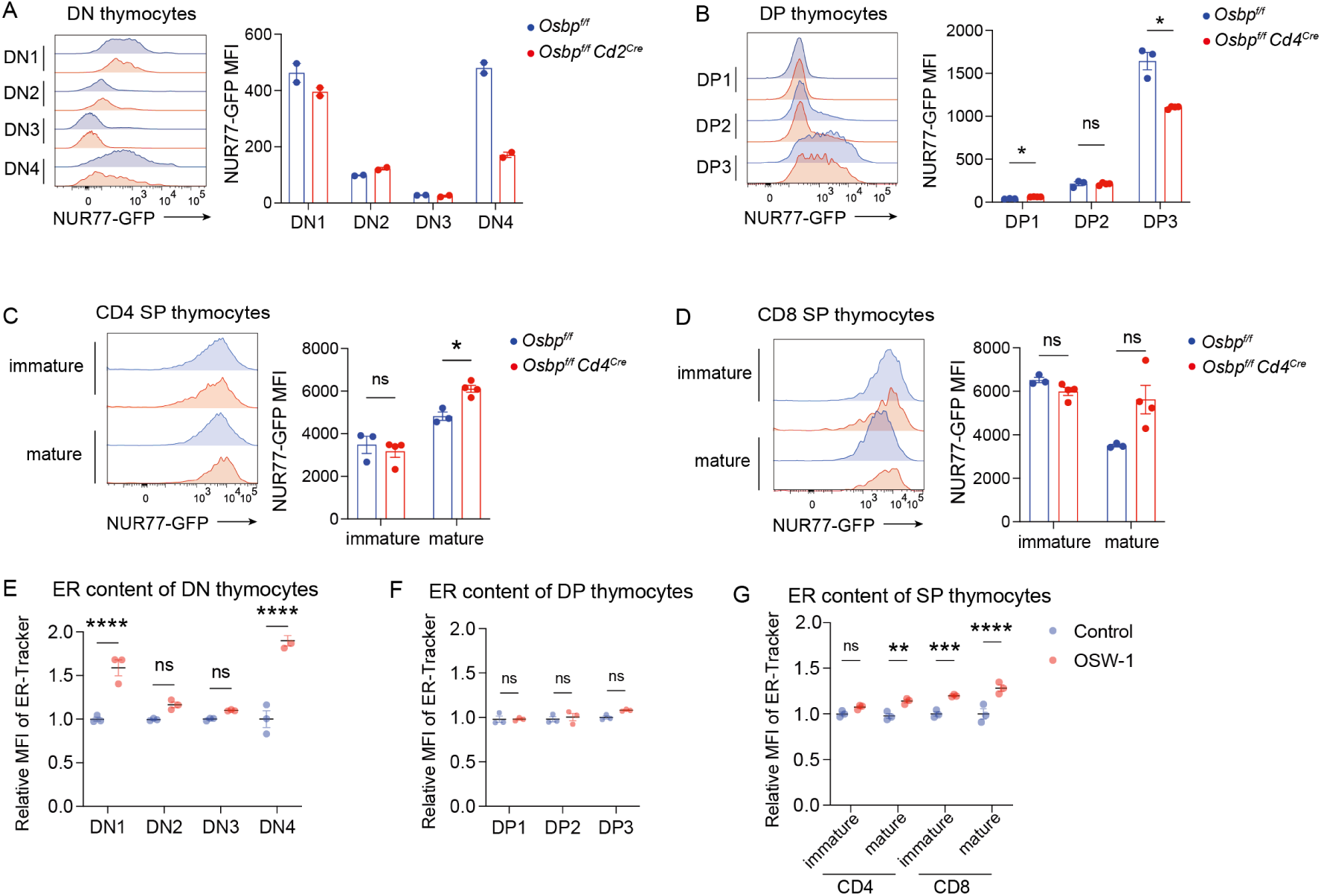
OSBP disruption impairs TCR signaling and ER homeostasis across thymocyte subsets. (A-D) NUR77-GFP reporter expression in the indicated thymocyte subsets. (A) NUR77-GFP MFI was quantified in DN1-DN4 thymocytes from *Osbp^f/f^ Cd2^Cre^* and *Osbp^f/f^*control mice (n = 2 mice), and in DP1-DP3 (B), CD4 SP (C), and CD8 SP (D) thymocytes from *Osbp^f/f^ Cd4^Cre^* and *Osbp^f/f^* control mice (n = 3 for *Osbp^f/f^* mice and n = 4 for *Osbp^f/f^ Cd4^Cre^* mice). (E-G) ER content in DN1-DN4 (E), DP1-DP3 (F), CD4 SP and CD8 SP (G) thymocyte populations following OSBP inhibition, assessed by ER-Tracker staining. Thymocytes were treated with OSW-1 (0.2 nM) or DMSO for 2 h, and ER-Tracker MFI was normalized to DMSO-treated controls within each subset (n = 3). Data are presented as mean ± s.e.m. Statistical significance was assessed by multiple unpaired two-tailed t tests (A-D) or two-way ANOVA (E-G). ns, not significant; *p < 0.05, **p < 0.01, ***p < 0.001, ****p < 0.0001.

In addition to signaling, we examined the effect of OSBP inhibition on ER homeostasis across different thymocyte subsets. To capture early cellular consequences, we used OSW-1 to exert acute OSBP inhibition and evaluated ER perturbation by ER-Tracker staining. Among DN subsets, DN1 and DN4 thymocytes exhibited the most pronounced ER perturbation following OSBP inhibition (Fig. 4E). Notably, DN4 thymocytes, which showed the strongest defects in pre-TCR signaling and developmental progression upon *Osbp* deficiency, also exhibited marked ER perturbation following acute OSBP inhibition. In contrast, DP thymocytes displayed comparatively modest ER changes after OSBP inhibition (Fig. 4F). Among SP thymocytes, CD8 SP cells exhibited greater ER perturbation than CD4 SP cells, with this difference becoming more evident during maturation (Fig. 4G).

Together, these findings reveal stage-specific patterns of OSBP vulnerability, explaining different phenotypes across developmental checkpoints. DN thymocytes exhibited defects in both pre-TCR signaling and ER homeostasis. DP thymocytes showed impaired TCR signaling while relatively preserved ER homeostasis. In contrast, SP thymocytes displayed greater disruption of ER homeostasis while preserving TCR signaling.

### Peripheral T-cell activation depends on OSBP

After establishing the essential role of OSBP in thymocyte development, we next examined its function in peripheral T cells. Because OSBP-deficient mice contained very few peripheral T cells, we used acute pharmacological inhibition to examine OSBP function in mature T cells. Resting T cells maintained relatively low PM cholesterol levels, which were only modestly affected by OSBP inhibition (Fig. 5A). Upon activation, however, both accessible PM cholesterol and total cellular cholesterol increased markedly, with CD8⁺ T cells exhibiting substantially higher levels than CD4⁺ T cells. OSBP inhibition caused only minor changes in total cholesterol but markedly reduced accessible PM cholesterol, with a more pronounced effect in CD8⁺ than in CD4⁺ T cells (Figs. 5A, B). Consistent with these changes, OSBP inhibition impaired TCR signaling, as indicated by reduced ERK phosphorylation, in both CD4⁺ and CD8⁺ T cells, with CD8⁺ T cells showing a greater reduction (Figs. 5C-E). In parallel, we assessed ER homeostasis and found that OSBP inhibition induced greater ER perturbation in activated CD8⁺ T cells than in activated CD4⁺ T cells (Fig. 5F). Notably, OSBP inhibition caused severe cell death in activated T cells, associated with profound phosphatidylserine exposure (Figs. 5G-H). These findings show that peripheral T cells exhibit increased cholesterol demands upon activation, rendering them increasingly dependent on OSBP-mediated cholesterol transport.

**Figure 5.**
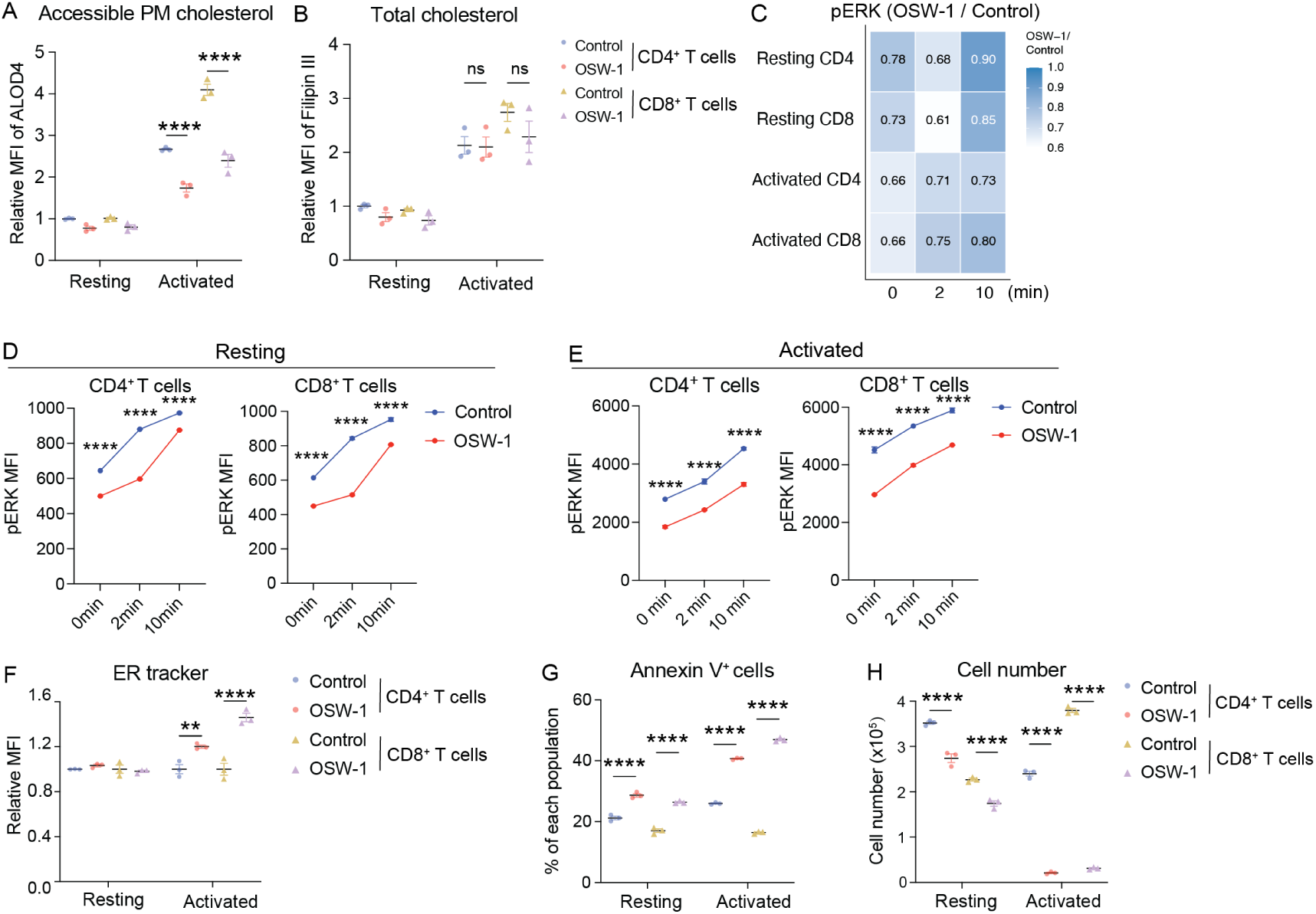
Peripheral T-cell activation increases dependence on OSBP-mediated cholesterol transport. (A-B) Accessible PM cholesterol (A) and total cellular cholesterol levels (B) in resting and activated peripheral CD4⁺ and CD8⁺ T cells following OSBP inhibition. T cells were treated with OSW-1 (0.2 nM, unless otherwise described) or DMSO for 2 h. ALOD4 and Filipin III staining were used to assess accessible PM cholesterol and total cellular cholesterol, respectively, with MFI values normalized to DMSO-treated resting CD4⁺ T cells (n = 4). (C-E) Effects of OSBP inhibition on TCR signaling in peripheral CD4⁺ and CD8⁺ T cells. Resting and activated T cells were pretreated with OSW-1 or DMSO control for 2 h and stimulated for the indicated times, followed by flow cytometric analysis of phosphorylated ERK (pERK). (C) Heatmap showing relative pERK MFI in resting and activated CD4⁺ and CD8⁺ T cells, normalized to the DMSO-treated control for each subset and time point. (D-E) Time-course analysis of pERK in resting (D) and activated (E) CD4⁺ and CD8⁺ T cells (n = 3). (F) ER content in resting and activated peripheral CD4⁺ and CD8⁺ T cells, assessed by ER-Tracker staining. Cells were treated with OSW-1 or DMSO for 2 h, and ER-Tracker MFI was normalized to the DMSO-treated control for each activation state and T-cell subset (n = 3). (G-H) Effects of OSBP inhibition on the survival of resting and activated T cells. Equal numbers of cells were treated with OSW-1 or DMSO control. Percentages of Annexin V^+^ cells were determined after 6 h (G), whereas viable (PI^-^) cells numbers were assessed after 10 h (H) (n = 3). Data are presented as mean ± s.e.m. Statistical significance was determined by two-way ANOVA (A, B, F-H) or unpaired two-tailed t test at individual time points (D, E). ns, not significant; **p < 0.01, ****p < 0.0001.

### Oxysterols perturb OSBP-mediated cholesterol transport in T cells

To further validate the role of OSBP in human T-cell function, we performed *OSBP* knockdown in human primary T cells. Similar to our findings in mouse T cells, *OSBP* depletion impaired TCR-induced phosphorylation of CD3ζ (pY142) and increased Annexin V positivity (Figs. 6A, B). Functionally, *OSBP*-depleted human T cells exhibited reduced cytotoxic activity against tumor cells (Fig. 6C).

**Figure 6.**
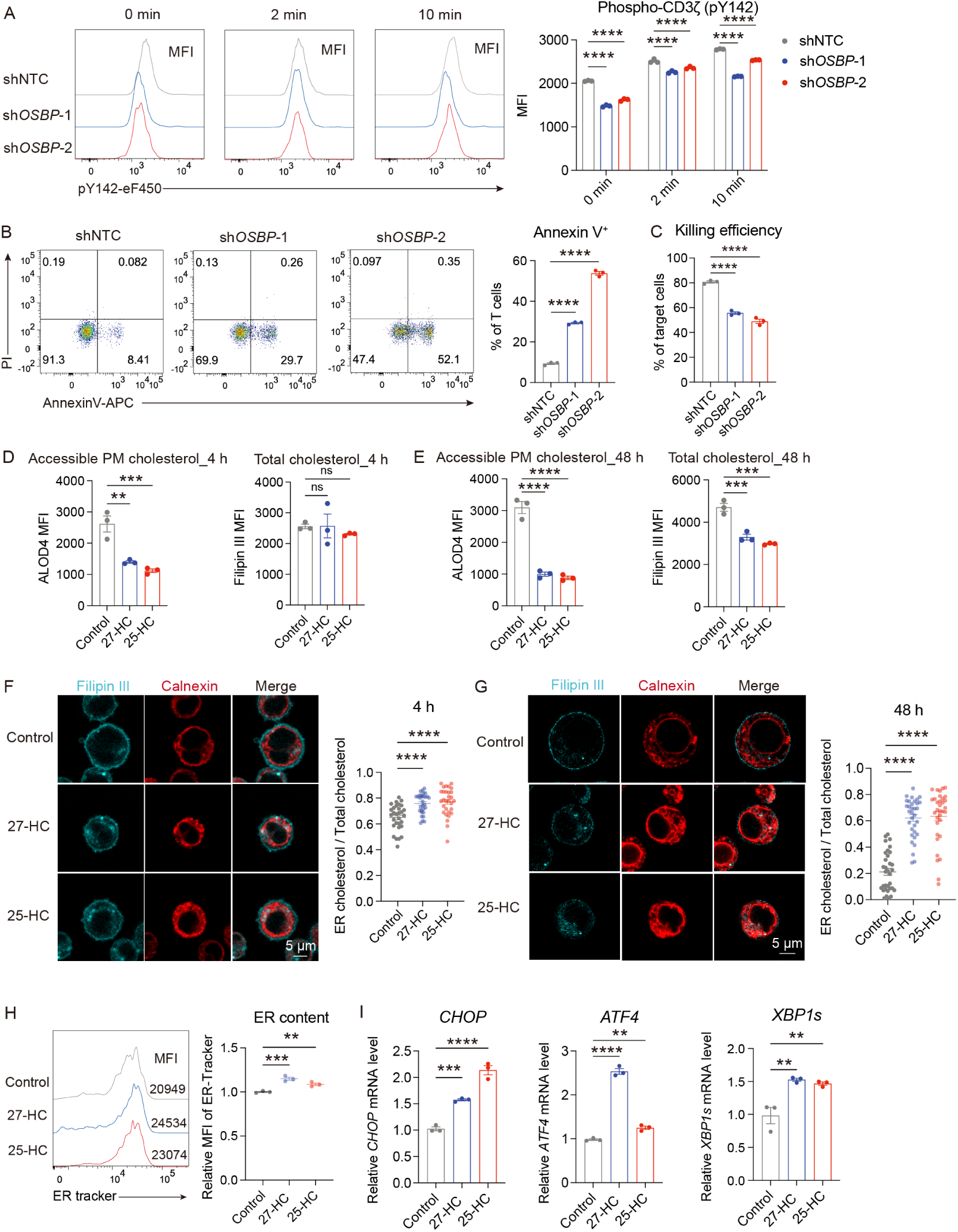
Oxysterols disrupt OSBP-mediated cholesterol transport in T cells. (A) Effects of *OSBP* knockdown on TCR signaling in human primary T cells. Following *OSBP* knockdown, cells were stimulated for the indicated times, and phospho-CD3ζ (pY142) was assessed by flow cytometry. Representative flow cytometry profiles and pCD3ζ MFI are shown (n = 3). (B) Annexin V/PI staining of control and *OSBP*-knockdown human primary T cells, with representative flow cytometry profiles and quantification of Annexin V^+^ cells (n = 3). (C) Cytotoxicity of human primary T cells following *OSBP* knockdown. Control and *OSBP*-knockdown T cells were cocultured with Raji cells at an effector-to-target ratio of 1:1 for 72 h. Cytotoxicity was assessed by the percentage of Raji cell killing, calculated based on the reduction in viable Raji cells after co-culture with T cells relative to Raji-only controls (n = 3). (D-E) Accessible PM cholesterol and total cellular cholesterol levels in human primary T cells following oxysterol treatment for 4 h (D) or 48 h (E), assessed by ALOD4 and Filipin III staining, respectively. PBMCs were stimulated with anti-CD3/CD28 beads for 3 days prior to treatment with MβCD-27-HC or MβCD-25-HC (1 μg/mL, unless otherwise described) (n = 3). (F-G) Subcellular cholesterol distribution in human primary T cells following MβCD-27-HC or MβCD-25-HC treatment for 4 h (F) or 48 h (G), under the conditions described in (D-E). The fraction of Filipin III signal overlapping with calnexin was quantified from confocal images using the thresholded Manders’ coefficient (n = 31 cells in F; n = 33 cells in G). Scale bar, 5 μm. (H) ER content in human primary T cells treated with MβCD-27-HC or MβCD-25-HC for 24 h. ER-Tracker MFI was normalized to MβCD-treated controls (n = 3). (I) Expression of ER stress-related genes following oxysterol treatment. Cells were treated with MβCD-27-HC or MβCD-25-HC for 48 h, and relative mRNA levels of *CHOP*, *ATF4*, and *XBP1s* were analyzed by quantitative PCR. Expression levels were normalized to MβCD-treated controls (n = 3). Data are presented as mean ± s.e.m. Statistical significance was assessed using two-way ANOVA (A) or one-way ANOVA (B-I). ns, not significant; **p < 0.01, ***p < 0.001, ****p < 0.0001.

Oxysterols are endogenous cholesterol metabolites that bind OSBP and modulate its cholesterol transport activity^43–45^. Previous studies have shown that oxysterols, particularly 27-hydroxycholesterol (27-HC) and 25-hydroxycholesterol (25-HC), accumulate in the tumor microenvironment and suppress T-cell immunity^3,46^. Short-term exposure (4 h) to either 27-HC or 25-HC selectively reduced accessible PM cholesterol while leaving total cellular cholesterol largely unchanged (Fig. 6D). In contrast, prolonged treatment (48 h) decreased both accessible PM cholesterol and total cellular cholesterol (Fig. 6E), consistent with the established role of oxysterols in limiting cellular cholesterol accumulation^47,48^. Importantly, both acute and prolonged oxysterol treatment increased ER cholesterol accumulation (Figs. 6F, G), recapitulating the cholesterol redistribution observed following OSBP inhibition. This redistribution was accompanied by disruption of ER homeostasis, as evidenced by increased ER content and elevated expression of ER stress-associated genes (Figs. 6H, I).

These findings identify oxysterols as endogenous inhibitors of OSBP-mediated cholesterol transport in T cells. By inhibiting cholesterol transport, oxysterols simultaneously reduce accessible PM cholesterol and induce ER cholesterol overload, providing a mechanistic explanation for the concurrent PM cholesterol deficiency and ER cholesterol accumulation previously observed in tumor-infiltrating T cells^3,49^.

## DISCUSSION

Here, we identify OSBP-mediated intracellular cholesterol transport as an essential determinant of T-cell development and activation. By coordinating cholesterol partitioning between the plasma membrane and the endoplasmic reticulum, OSBP simultaneously sustains TCR signaling competence and preserves ER homeostasis. OSBP deficiency causes profound defects at multiple developmental checkpoints, including β-selection, positive selection, and post-positive-selection maturation. In peripheral T cells, OSBP is indispensable for activation, and its disruption results in extensive cell death. Moreover, oxysterols act as endogenous inhibitors of OSBP, recapitulating the disruption of spatial cholesterol homeostasis and associated T-cell dysfunction (Fig. S4).

The plasma membrane is highly enriched in cholesterol, which supports both membrane signaling and mechanical stability^18,50,51^. Cholesterol promotes TCR clustering and downstream phosphorylation events, as well as the assembly of signaling nanodomains^23,52^, thereby providing a mechanistic basis for the impaired TCR signaling observed following OSBP disruption. In contrast, the ER contains only a small fraction of cellular cholesterol and has a relatively thin, highly disordered membrane enriched in unsaturated phospholipids^53^. These biophysical properties are well suited to support protein synthesis, membrane protein insertion, quality control and secretory trafficking. Consistent with this framework, ER cholesterol accumulation caused by OSBP disruption robustly activates the unfolded protein response and ultimately leads to extensive cell death. These findings highlight that intracellular cholesterol transport is required to preserve the distinct biophysical identities of the PM and ER, thereby supporting both receptor signaling and ER fitness.

The requirement for OSBP is closely coupled to the activity of cholesterol biosynthesis. TCR and pre-TCR signaling trigger mTORC activation^54^, leading to robust cholesterol biosynthesis in the ER. Under these conditions, OSBP is required to efficiently transport newly synthesized cholesterol to the PM, where it supports immunoreceptor signaling while preventing cholesterol overload in the ER. Accordingly, the effects of OSBP disruption were most pronounced in DN thymocytes and activated T cells, which exhibited the highest cholesterol levels among the T-cell populations examined. It is interesting to note that blocking cholesterol biosynthesis yields only minor defects in thymocyte development^3,55^. Similarly, manipulation of cholesterol abundance through cholesterol uptake^56^, efflux^57,58^, or esterification^12^ has limited effects on thymocyte development. Together, these findings indicate that the intracellular distribution of cholesterol, rather than its total abundance alone, is indispensable for normal thymocyte development.

It is informative to compare the distinct roles of anterograde and retrograde PM cholesterol transport pathways during thymocyte development. Aster-A mediates retrograde cholesterol transport from the PM to the ER^21^. Although Aster-A deficiency alters PM cholesterol organization and TCR signaling, its impact on thymocyte development is limited. In contrast, OSBP deficiency simultaneously disrupts PM signaling and induces severe ER stress through ER cholesterol accumulation. Given that developing thymocytes are particularly sensitive to ER stress^59,60^, the combined impairment of membrane signaling and ER homeostasis likely accounts for the profound developmental defects caused by OSBP loss.

Our findings also uncover a previously unrecognized mechanism underlying the immunomodulatory effects of oxysterols. Oxysterols accumulate in diverse pathological settings, such as cancer and obesity^3,61–63^. They are classically viewed as regulators of cholesterol metabolism through inhibition of SREBP2 and activation of LXR^48,64^, thereby suppressing cholesterol synthesis and uptake while promoting cholesterol efflux. These transcriptional responses typically require hours to days to affect cellular cholesterol abundance. In contrast, we show that acute oxysterol exposure rapidly inhibits OSBP-mediated cholesterol transport without measurably altering total cellular cholesterol levels, resulting in a marked reduction in accessible PM cholesterol and concomitant cholesterol accumulation in the ER. Thus, disruption of spatial cholesterol homeostasis represents a rapid, non-transcriptional mechanism by which oxysterols suppress T-cell function. This mechanism may help explain previous observations that tumor-infiltrating T cells display both reduced PM cholesterol availability and elevated ER cholesterol^3,49^. Likewise, obesity is associated with increased circulating and tissue oxysterol levels, impaired thymopoiesis, and enhanced thymocyte apoptosis^65,66^, raising the possibility that pathological oxysterol accumulation compromises T-cell development through inhibition of OSBP-dependent cholesterol transport.

Together, our study establishes OSBP-mediated spatial cholesterol homeostasis as a fundamental metabolic determinant of T-cell development and activation. More broadly, our findings expand the paradigm of cholesterol regulation in T cells beyond total cholesterol abundance to encompass its intracellular distribution, and suggest a potential therapeutic avenue for modulating immune responses in both physiological and pathological settings.

## MATERIALS AND METHODS

### Mice

*Osbp^flox/flox^* mice, in which exon 2 of the *Osbp* gene is flanked by loxP sites, were generated by Yingjie Wu (Dalian Medical University, China). *Orp2^flox/flox^* mice were provided by Dr. Mingming Gao (Hebei Medical University, China). The *hCD2^iCre^* (referred to as *Cd2^Cre^*) and *Cd4^Cre^* transgenic mice were provided by Xiaolong Liu (Shanghai Institute of Biochemistry and Cell Biology, China), and NUR77-GFP transgenic mice were provided by Haopeng Wang (ShanghaiTech University, China). C57BL/6J CD45.2 wild-type mice were obtained from LingChang (Shanghai, China). OT-I TCR transgenic mice (RRID: IMSR_JAX:003831) were obtained from The Jackson Laboratory. T-cell-specific *Osbp*-deficient mice were generated by crossing *Osbp^f/f^* mice with *Cd2^Cre^* or *Cd4^Cre^* mice. Age-matched littermates aged 3-4 weeks were used for developmental analyses, whereas mice aged 6-8 weeks were used for all other experiments.

All mice were maintained under specific pathogen-free conditions in the animal facility of the Shanghai Institute of Biochemistry and Cell Biology. All animal procedures were approved by the Institutional Animal Care and Use Committee of the Shanghai Institute of Biochemistry and Cell Biology, Chinese Academy of Sciences (ethical approval no. SIBCB-S331-2306-22), and conducted in accordance with institutional and national guidelines.

### Cell lines

HEK-293FT cells were purchased from the Cell Bank of the Chinese Academy of Sciences (Shanghai, China) and cultured in Dulbecco’s modified Eagle medium (DMEM) supplemented with 10% fetal bovine serum (FBS), 100 U/mL penicillin, and 100 μg/mL streptomycin. Jurkat T cells (RRID: CVCL_0367; male) and Raji cells (RRID: CVCL_0511; male) were cultured in RPMI 1640 medium supplemented with 10% FBS, 100 U/mL penicillin, and 100 μg/mL streptomycin.

### Primary cell cultures

Primary mouse T cells were isolated from the spleen and cultured in RPMI 1640 medium supplemented with 10% FBS, 100 U/mL penicillin, 100 μg/mL streptomycin, 50 μM 2-mercaptoethanol, and recombinant IL-2 (Peprotech, Cat. 200-02, 10 ng/mL). Human peripheral blood mononuclear cells (PBMCs) were isolated from healthy donors by density-gradient centrifugation and cryopreserved in liquid nitrogen until use. PBMCs were cultured in X-VIVO 15 medium supplemented with 5% FBS, 100 U/mL penicillin, 100 μg/mL streptomycin, 10 mM N-acetyl-L-cysteine, and 10 ng/mL IL-2. All cells were incubated at 37 °C in a humidified atmosphere containing 5% CO₂ and were routinely tested and confirmed to be negative for mycoplasma contamination.

### Lentiviral transduction

Lentivirus was produced by transfecting HEK-293FT cells with the packaging plasmid psPAX2 and the envelope plasmid pMD2.G, together with pHAGE-based overexpression or pLKO.1-based knockdown vectors using Lipofectamine 2000 (Invitrogen, Cat. 11668019) according to the manufacturer’s instructions. Virus-containing supernatants were harvested 48 h after transfection. Jurkat T cells were transduced with viral supernatants for 24 h, after which the supernatants were removed. Cells were further cultured and sorted by fluorescence-activated cell sorting (FACS) before downstream analyses.

For human primary T-cell transduction, PBMCs were activated with Human T-Activator CD3/CD28 Dynabeads (Gibco, Cat. 11132D) for 24 h. Viral particles were concentrated overnight from the supernatants using 8% PEG8000, pelleted by centrifugation, and resuspended in fresh medium. Activated cells were subsequently transduced with the concentrated virus for 48 h, after which the virus-containing medium and Dynabeads were removed. The transduced cells were further cultured and sorted by FACS before downstream assays.

### Flow cytometry

For surface marker staining, cells were incubated with fluorophore-conjugated antibodies at 4 °C for 30 min and washed prior to analysis or subsequent staining. For intracellular staining, surface-stained cells were fixed with eBioscience IC Fixation Buffer (Invitrogen, Cat. 00-8222-49), permeabilized with eBioscience Permeabilization Buffer (Invitrogen, Cat. 00-8333-56), and stained with antibodies against intracellular targets according to the manufacturer’s instructions. For analysis of ER content, cells were incubated with ER-Tracker (Invitrogen, Cat. E34251, 1 μM) at 37 °C for 20 min and, where required, subsequently stained with surface markers to distinguish T-cell subsets. Flow cytometry data were acquired using an Attune NxT flow cytometer (Thermo Fisher Scientific) and analyzed with FlowJo software. Antibodies used for flow cytometry are listed in Supplementary Table 1.

### Measurement of cholesterol levels

#### ALOD4 staining for accessible plasma membrane cholesterol

Accessible PM cholesterol was quantified using the cholesterol-binding probe ALOD4. Recombinant ALOD4-mCherry was expressed in *E. coli* and purified by affinity chromatography, and Alexa Fluor 647 (Invitrogen, Cat. A20347)-labeled ALOD4 was prepared according to a published protocol^20^. Cells were resuspended at 2 × 10⁶ cells/mL in serum-free DMEM and incubated with ALOD4 (20 μg/mL) at 37 °C for 30 min, followed by surface marker staining at 4 °C for 30 min and fixation and permeabilization for intracellular staining when required.

#### Measurement of total cellular cholesterol

For flow cytometric analysis of total cellular cholesterol using Filipin III staining, cells were fixed with IC Fixation Buffer and stained with Filipin III (Cayman Chemical, Cat. 70440, 50 μg/mL) at 4 °C overnight. For confocal microscopy, cells were allowed to adhere to poly-L-lysine-coated dishes, fixed, and stained with Filipin III (50 μg/mL) at 4 °C overnight.

For biochemical quantification of total cellular cholesterol, cellular lipids were extracted by resuspending cells in 0.1 mL ddH₂O followed by addition of 0.4 mL methanol/chloroform (1:2, v/v). The lower organic phase was collected and dried, and total cholesterol was quantified using the Amplex Red Cholesterol Assay Kit (Invitrogen, Cat. A12216) according to the manufacturer’s instructions. Cholesterol levels were normalized to cell number.

#### Thymocyte isolation and developmental analysis

Thymocytes were isolated from thymic lobes by gentle mechanical dissociation through a 40-μm cell strainer, followed by red blood cell lysis and staining with fluorophore-conjugated antibodies for flow cytometric analysis.

For developmental profiling, thymocytes were classified into double-negative (DN; CD4^-^CD8^-^), double-positive (DP; CD4^+^CD8^+^), and single-positive (SP, CD4^+^CD8^-^ or CD4^-^CD8^+^) populations. DN cells were further subdivided into DN1 (CD44^+^CD25^-^), DN2 (CD44^+^CD25^+^), DN3 (CD44^-^CD25^+^), and DN4 (CD44^-^CD25^-^) populations; DP cells were subdivided into DP1 (CD69^lo^ TCRβ^lo^), DP2 (CD69^hi^ TCRβ^mid^), and DP3 (CD69^hi^ TCRβ^hi^) populations, whereas TCRβ^+^ SP thymocytes were classified as immature (CD24^+^) or mature (CD24^-^).

#### TCR signaling assays

Mouse T cells were first isolated from mouse spleens by negative selection using a Mouse T Cell Isolation Kit (StemCell Technologies, Cat. 19851A) and allocated to resting or activated conditions. Resting T cells were maintained without stimulation, whereas activated T cells were generated by stimulation with plate-bound anti-mCD3ε (Clone 145-2C11, BioLegend, Cat. 100340, 5 μg/mL) and anti-mCD28 (Clone 37.51, BioLegend, Cat. 102116, 5 μg/mL) for 2 days, followed by expansion for 5 days in the medium described above. For phospho-flow cytometric analysis of mouse peripheral T cells, cells were suspended in serum-free DMEM at a density of 5 × 10^6^ cells/mL and stimulated with biotinylated anti-mCD3ε (Clone 145-2C11, BioLegend, Cat. 100304, 5 μg/mL) and anti-mCD28 (Clone 37.51, BioLegend, Cat. 102104, 5 μg/mL) for the indicated times. The stimulation antibodies were pre-crosslinked with streptavidin (Invitrogen, Cat. 434301, 5 μg/mL). Cells were subsequently stained for surface markers on ice, followed by intracellular staining for phospho-ERK1/2.

For measurement of TCR signaling in Jurkat cells, the treated cells were suspended in serum-free RPMI-1640 medium at a density of 5 × 10^7^ cells/mL and stimulated with anti-hCD3ε (Clone OKT3, BioLegend, Cat. 317302, 10 μg/mL) pre-crosslinked with anti-mouse IgG (Invitrogen, Cat. 31160, 10 μg/mL) at 37 °C. Cells were immediately lysed in RIPA buffer supplemented with protease and phosphatase inhibitors at the indicated time points. Phosphorylated and total CD3ζ, ZAP70, PLCγ1, and ERK levels were analyzed by immunoblotting using antibodies listed in Supplementary Table 1.

For human primary T cells, cells were suspended in serum-free DMEM at a density of 5 × 10⁶ cells/mL and stimulated with anti-hCD3ε (Clone OKT3, BioLegend, Cat. 317302, 5 μg/mL) and anti-hCD28 (Clone CD28.2, BioLegend, Cat. 302933, 5 μg/mL) pre-crosslinked with anti-mouse IgG (5 μg/mL) for the indicated times. Cells were subsequently fixed, permeabilized, and stained intracellularly for phospho-CD3ζ (Y142), followed by flow cytometric analysis.

#### Cell number analysis following OSW-1 treatment

For Jurkat cell number analysis, cells were seeded at 5 × 10⁵ cells/mL and treated with OSW-1 (MCE, HY-101213, 5 nM) or DMSO for 20 h. Absolute cell numbers were determined by flow cytometry. For mouse T-cell analysis, resting T cells were maintained without stimulation, whereas activated T cells were generated under the conditions described above. Cells were resuspended at a density of 1 × 10⁶ cells/mL and subsequently treated with OSW-1 (0.2 nM) or DMSO for 10 h. CD4⁺ and CD8⁺ T cells were identified by surface staining, and absolute cell numbers were determined by flow cytometry.

#### Sterol treatment of Jurkat and human primary T cells

Cholesterol (Sigma-Aldrich, Cat. C8667), 25-hydroxycholesterol (Avanti Polar Lipids, Cat. 700019P), or 27-hydroxycholesterol (Avanti Polar Lipids, Cat. 700021P) was separately loaded onto methyl-β-cyclodextrin (MβCD, Sigma-Aldrich, Cat. 332615) according to a previously described method^25^. For cholesterol replenishment, Jurkat T cells were treated with MβCD-cholesterol (5 μg/mL) for 4 h at 37 °C. For oxysterol treatment, human primary T cells activated with CD3/CD28 Dynabeads for 3 days were treated with MβCD-25-HC or MβCD-27-HC at concentrations and for durations specified in the corresponding figure legends.

#### T cell cytotoxicity assay

Raji cells were labeled with CellTracker-Violet (Invitrogen, Cat. C34557) and cocultured with control or *OSBP*-knockdown activated human primary T cells at an effector-to-target ratio of 1:1 in 96-well plates for 72 h. Cells were subsequently stained with propidium iodide (PI), and viable Raji cells were identified as CellTracker^+^ PI^-^cells by flow cytometry. Cytotoxic activity was calculated based on the reduction in viable Raji cell numbers in coculture relative to Raji-only controls as follows: killing efficiency (%) = [1 − (viable Raji cells in coculture/viable Raji cells in Raji-only controls)] × 100.

#### Confocal imaging of ER-associated cholesterol

Cells were seeded onto poly-L-lysine-coated dishes, fixed, and stained with Filipin III and anti-calnexin (Abcam, Cat. ab22595) overnight at 4 °C, followed by anti-rabbit IgG-AF647 (BioLegend, Cat. 406414) staining at room temperature for 1 h. Images were acquired using an Olympus FV3000 confocal microscope under identical settings and analyzed with Fiji. ER-associated cholesterol was quantified using the thresholded Manders’ coefficient (tM1), with Filipin III designated as channel 1 and calnexin as channel 2.

#### Transmission electron microscopy

Jurkat cells treated with OSW-1 (5 nM, 20 h) or vehicle control were fixed overnight at 4 °C in 2.5% glutaraldehyde in PBS (pH 7.4), postfixed with 1% osmium tetroxide, stained en bloc with 2% uranyl acetate, dehydrated through a graded ethanol series, and embedded in Spurr’s resin. Ultrathin sections were prepared using a Leica EM UC7 ultramicrotome, counterstained, and examined using a Tecnai G2 Spirit transmission electron microscope operated at 120 kV. Images were analyzed using Fiji.

#### Quantitative real-time PCR

Total RNA was extracted using TRIzol reagent (Invitrogen, Cat. 15596018) and reverse-transcribed into cDNA using the PrimeScript RT Reagent Kit with gDNA Eraser (Takara, Cat. RR047A). Quantitative real-time PCR was performed using SYBR Green Master Mix (Novoprotein, Cat. E096-01B), and relative gene expression was normalized to 18S rRNA and calculated using the 2^−ΔΔCt^ method. Primer sequences are listed in Supplementary Table 2.

#### Statistical analysis

Statistical analysis was performed using GraphPad Prism 10. Data are presented as mean ± s.e.m. Two-tailed t tests or one- or two-way ANOVA followed by appropriate multiple-comparisons tests were used as specified in the corresponding figure legends. Statistical significance was defined as \**p* < 0.05, \*\**p* < 0.01, \*\*\**p* < 0.001, and \*\*\*\**p* < 0.0001.

## DATA AVAILABILITY

All the data and materials related to this study are available from the corresponding author upon request.

## ACKNOWLEDGEMENTS

We thank the Core Facility of Chemical Biology and Molecular Biology, Cellular Biology, and Animal Facility at the CEMCS. This study is supported by the Ministry of Science and Technology of the Peoples’s Republic of China (MOST) (2023YFA1800200), the Strategic Priority Research Program of the Chinese Academy of Science (XDB0990000), C.X. is also a scholar of Shanghai Academy of Natural Sciences (SANS).

## AUTHOR CONTRIBUTIONS

C.X. and X.L. conceived the study. Y.L., X.H., C.L., and Z.R. designed and performed experiments. C.L. and Z.R. conducted the ORP family screen. X.H. and C.L. performed phenotypic analyses of *Osbp* and *Orp2* conditional knockout mice. Y.L. and X.H. carried out OSBP inhibition experiments. Y.L. performed experiments involving human primary T cells and oxysterol treatment experiments. Y.W., J.H., and Y.Q. generated the *Osbp^flox/flox^*mice, and M.G. provided the *Orp2^flox/flox^* mice. Y.L., X.H., C.X. and X.L. wrote the manuscript and other authors revised it.

---

**Figure S1.**
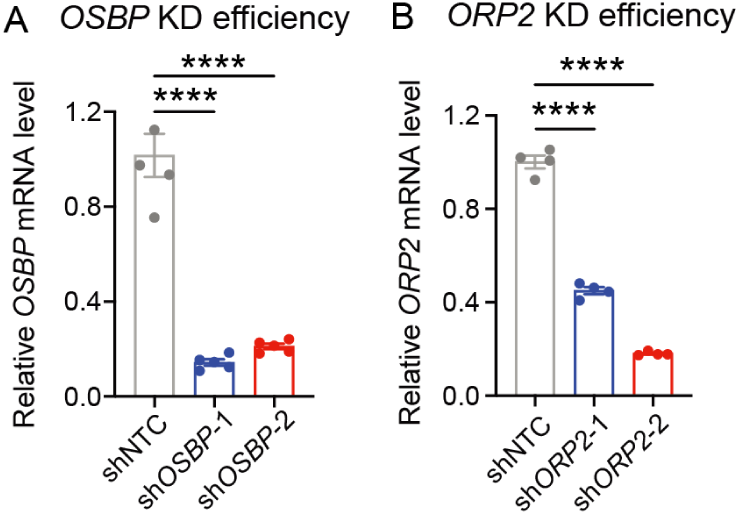
Validation of *OSBP* and *ORP2* knockdown efficiency in Jurkat T cells. (A-B) Relative *OSBP* (A) and *ORP2* (B) mRNA expression levels in Jurkat T cells following shRNA-mediated knockdown, determined by quantitative PCR. Values were normalized to those in cells transduced with non-targeting shRNA controls (n = 4). Data are presented as mean ± s.e.m. Statistical significance was determined by one-way ANOVA. ****p < 0.0001.

**Figure S2.**
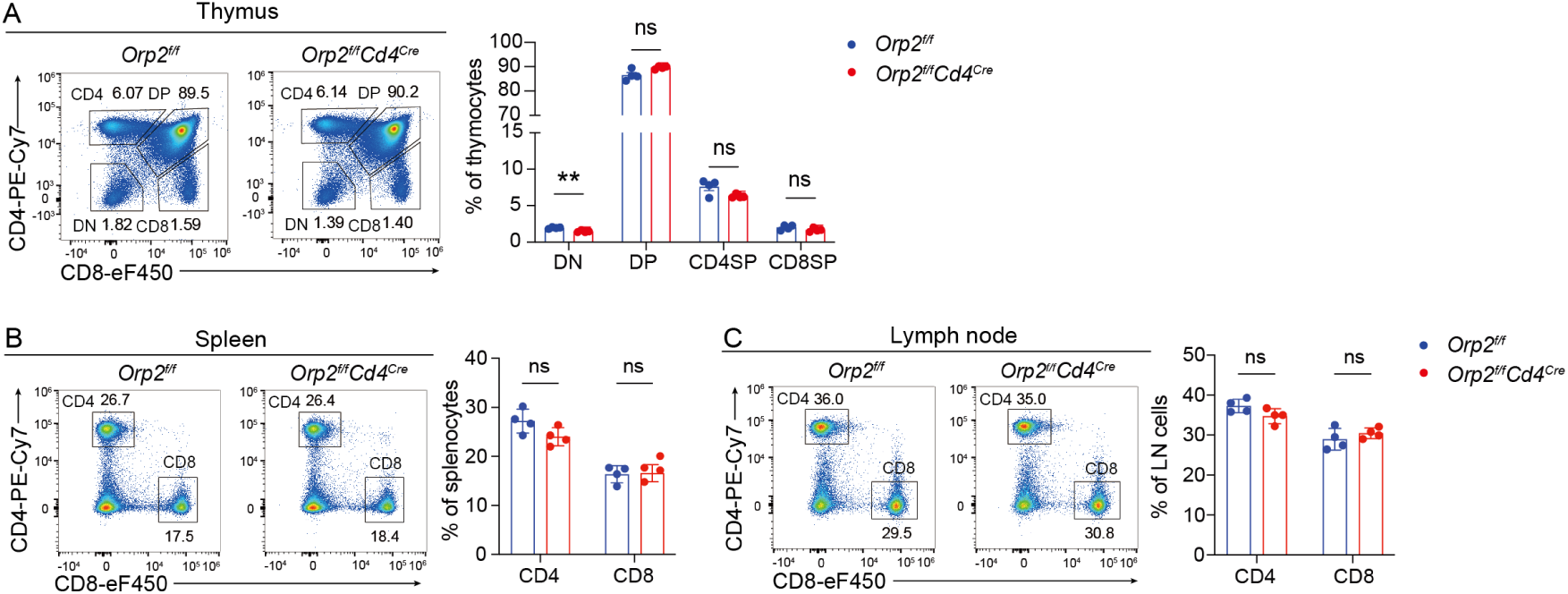
ORP2 deficiency does not affect thymocyte development or peripheral T-cell homeostasis. (A) Thymocyte development in *Orp2^f/f^ Cd4^Cre^* mice and *Orp2^f/f^* control mice. Representative flow cytometry profiles and percentages of DN, DP, CD4 SP, and CD8 SP populations among total thymocytes are shown (n = 4 mice). (B-C) Peripheral T-cell populations in the spleen (B) and inguinal lymph nodes (C) of *Orp2^f/f^ Cd4^Cre^* and *Orp2^f/f^* mice. Frequencies of CD4⁺ and CD8⁺ T cells among total cells were quantified (n = 4 mice). Data are presented as mean ± s.e.m. Statistical significance was assessed by multiple unpaired two-tailed t tests (A-C). ns, not significant.

**Figure S3.**
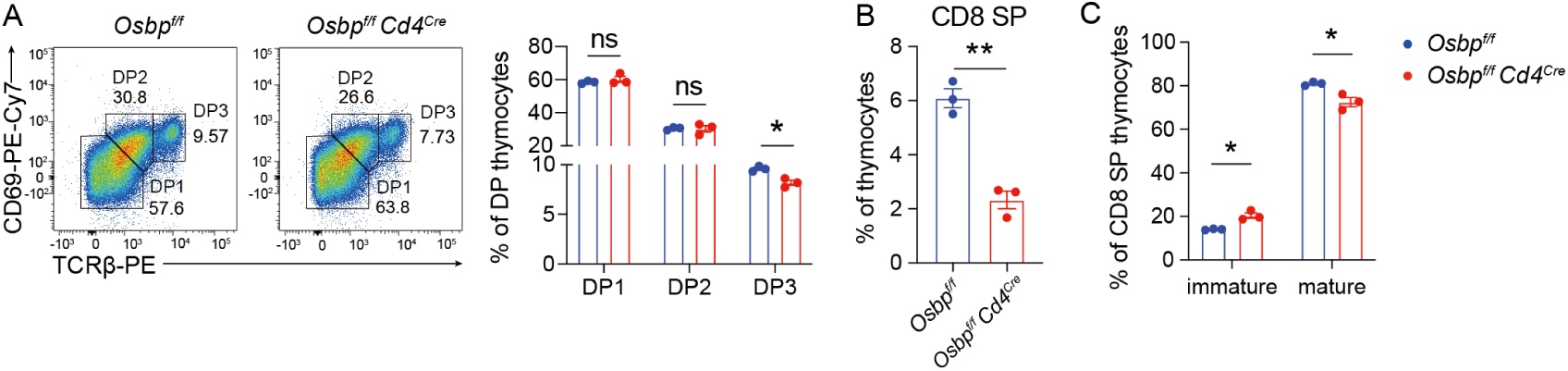
OSBP deficiency impairs positive selection and CD8 SP thymocyte maturation in OT-I TCR transgenic mice. (A-C) Thymocyte development in *Osbp^f/f^ Cd4^Cre^* OT-I and *Osbp^f/f^* OT-I control mice (n = 3 mice). (A) Frequencies of DP subsets within the DP population. (B) Frequencies of CD8 SP thymocytes among total thymocytes. (C) Percentages of immature and mature CD8 SP thymocytes. Data are presented as mean ± s.e.m. Statistical significance was determined using multiple unpaired two-tailed t tests (A, C) or an unpaired two-tailed t test (B). ns, not significant; *p < 0.05, **p < 0.01.

**Figure S4.**
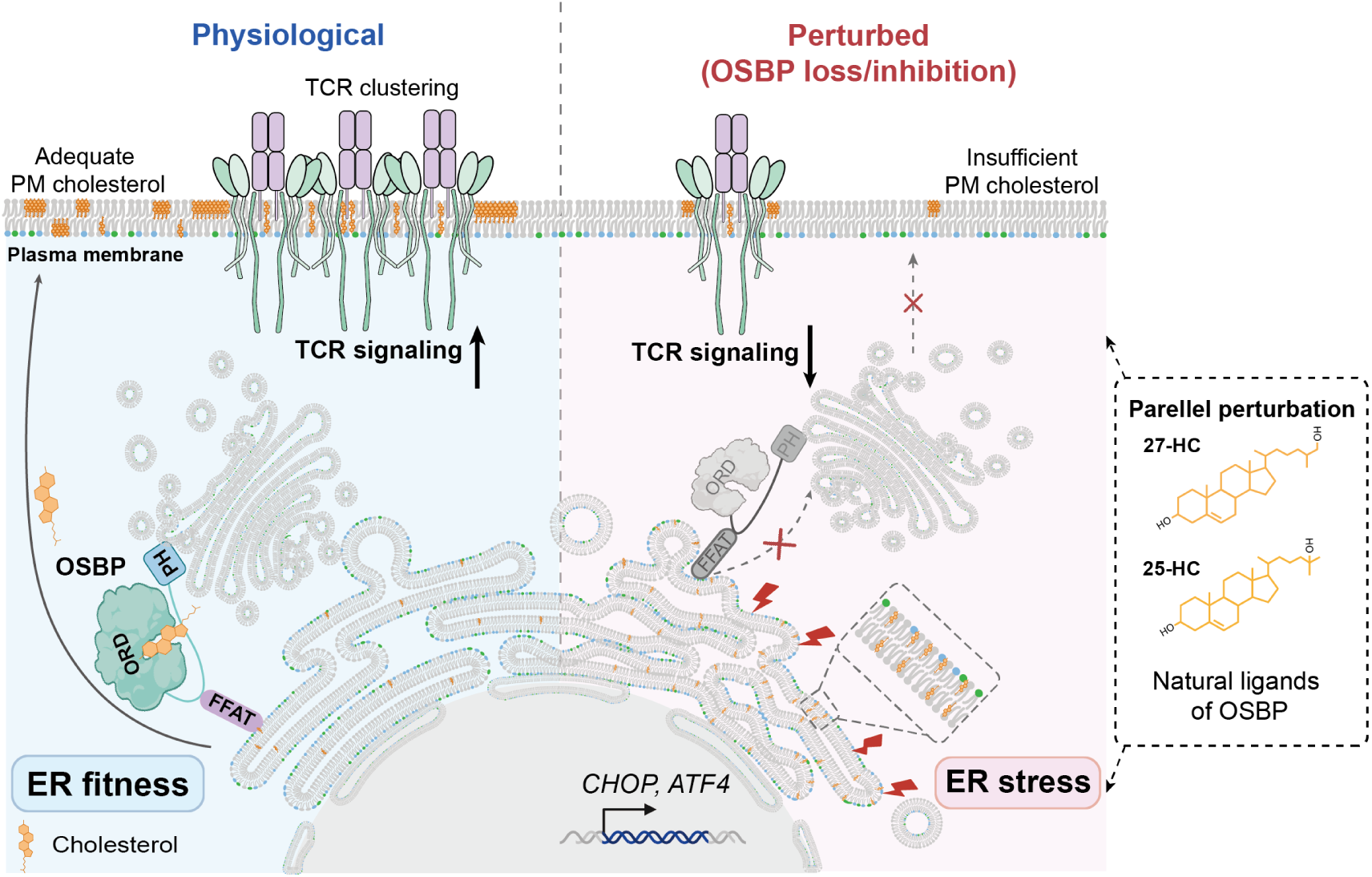
Schematic model summarizing the role of OSBP-mediated intracellular cholesterol transport in T cells. OSBP-mediated intracellular cholesterol transport maintains spatial cholesterol homeostasis in T cells by coordinating cholesterol partitioning between the plasma membrane and the endoplasmic reticulum. Through this coordination, OSBP maintains plasma membrane cholesterol levels to support TCR signaling competence while preventing ER cholesterol accumulation to preserve ER homeostasis and T-cell fitness. OSBP deficiency disrupts this balance, leading to impaired TCR signaling and induction of ER stress. Oxysterols act as natural OSBP inhibitors, resulting in altered cholesterol distribution and T-cell dysfunction.

**Supplementary Table 1.**

| ANTIBODIES | SOURCE | IDENTIFIER |
| --- | --- | --- |
| anti-mCD4-eFluor 450 | Invitrogen | 48-0041-82 |
| anti-mCD4-PE-Cy7 | Invitrogen | 25-0041-82 |
| anti-mCD4-BV786 | BD Biosciences | 563727 |
| anti-mCD4-BV750 | BD Biosciences | 747344 |
| anti-mCD8-PE | Invitrogen | 12-0081-82 |
| anti-mCD8-FITC | Invitrogen | 11-0081-81 |
| anti-mCD8-eFluor 450 | Invitrogen | 48-0081-82 |
| anti-mCD8-APC | Invitrogen | 17-0081-82 |
| anti-mCD44-PE-Cy7 | Invitrogen | 25-0441-82 |
| anti-mCD25-AF488 | Invitrogen | 53-0251-82 |
| anti-mCD25-APC | Invitrogen | 17-0251-82 |
| anti-mCD69-PE | Invitrogen | 12-0691-81 |
| anti-mCD69-PE-Cy7 | Invitrogen | 25-0691-82 |
| anti-mTCRβ-BV421 | BioLegend | 109230 |
| anti-mTCRβ-PE | Invitrogen | 12-5961-82 |
| anti-mCD24-BV421 | BioLegend | 101826 |
| anti-hCD69-PE | BioLegend | 310910 |
| anti-hCD45-AF647 | Invitrogen | 51-0459-42 |
| anti-pY142- eFluor 450 | Invitrogen | 48-2478-42 |
| donkey-anti-rabbit IgG-AF647 | Invitrogen | A31573 |
| Annexin V-APC | Invitrogen | 88-8007-74 |
| PI | Invitrogen | 00-6990-50 |
| LIVE/DEAD Fixable Near IR 780 | Invitrogen | L34994 |
| anti-CD16/CD32 | Invitrogen | 16-0161-85 |
| Rabbit polyclonal anti-Phospho-ZAP70 (Y319) | Cell Signaling | 2701S |
| Rabbit polyclonal anti-Phospho-PLCγ1 (Tyr783) | Cell Signaling | 2821S |
| Rabbit polyclonal anti-Phospho-ERK1/2 (Thr202/Tyr204) | Cell Signaling | 9101S |
| Rabbit monoclonal anti-pCD3ζ (Y142) | Abcam | ab68235 |
| Rabbit monoclonal anti-ZAP70 | Abcam | ab32410 |
| Rabbit polyclonal anti-PLCγ1 | Santa Cruze Biotechnology | sc-81 |
| Rabbit monoclonal anti-ERK1/2 | Cell Signaling | 4695S |
| Mouse monoclonal anti-CD3ζ | Santa Cruze Biotechnology | sc-1239 |
| GAPDH | KangChen Bio-tech | KC-5G5 |
| HRP linked anti-mouse IgG | Cell Signaling | 7076S |
| HRP linked anti-rabbit IgG | Cell Signaling | 7074S |

**Supplementary Table 2.**
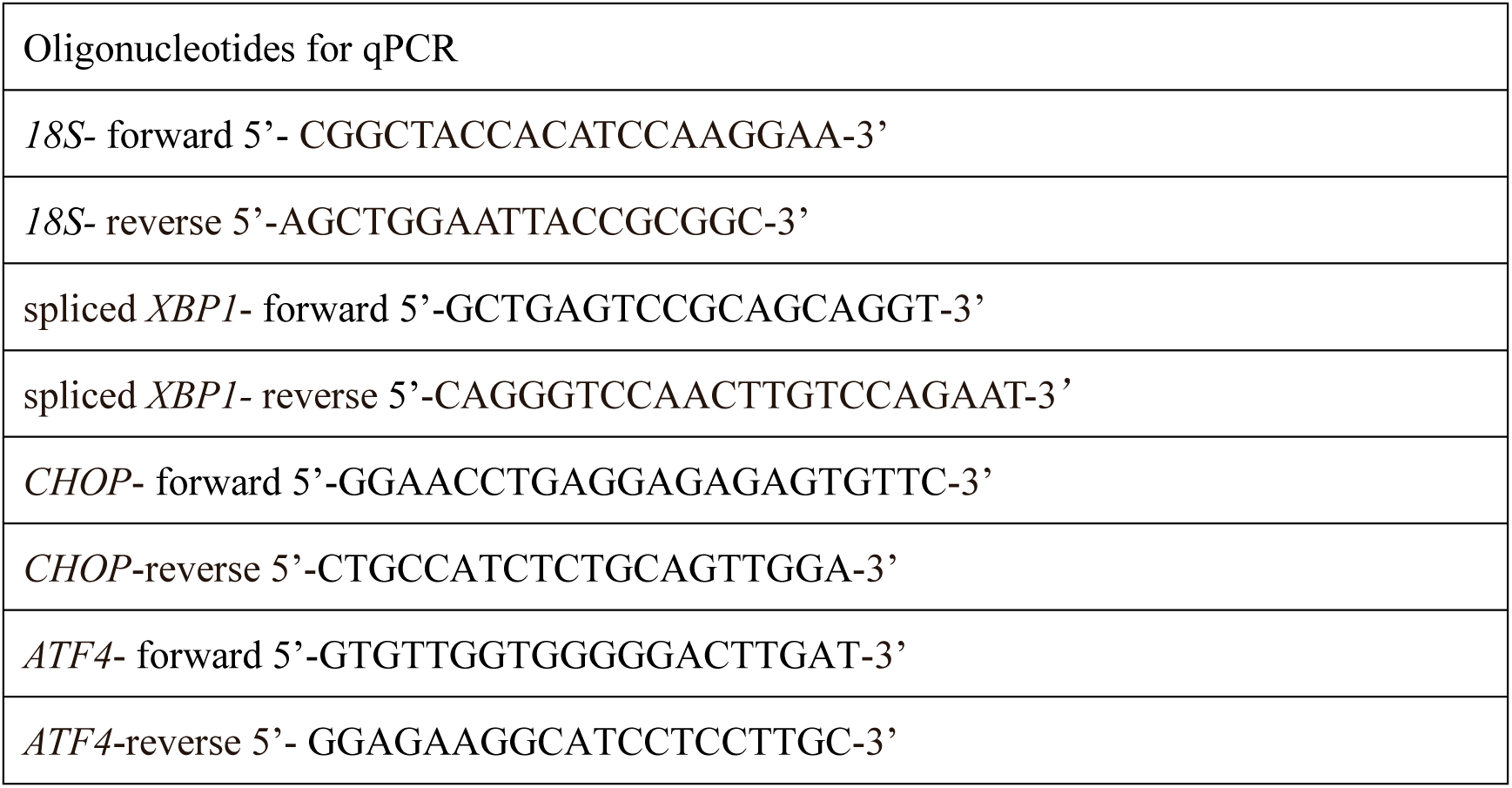

